# Thermal versus Mechanical Unfolding in a Model Protein

**DOI:** 10.1101/816801

**Authors:** Rafael Tapia-Rojo, Juan J. Mazo, Fernando Falo

## Abstract

Force spectroscopy techniques are often used to learn about the free energy landscape of single biomolecules, typically by recovering free energy quantities that, extrapolated to zero force, are compared to those measured in bulk experiments. However, it is not always clear how the information obtained from a mechanically perturbed system can be related to that obtained using other denaturants, since tensioned molecules unfold and refold along a reaction coordinate imposed by the force, which is unlikely meaningful in its absence. Here, we explore this dichotomy by investigating the unfolding landscape of a model protein, which is first unfolded mechanically through typical force spectroscopy-like protocols, and next thermally. When unfolded by non-equilibrium force extension and constant force protocols, we recover a simple two-barrier landscape, as the protein reaches the extended conformation through a metastable intermediate. Interestingly, folding-unfolding equilibrium simulations at low forces suggested a totally different scenario, where this metastable state plays little role in the unfolding mechanism, and the protein unfolds through two competing pathways^27^. Finally, we use Markov state models to describe the configurational space of the unperturbed protein close to the critical temperature. The thermal dynamics is well understood by a one-dimensional landscape along an appropriate reaction coordinate, however very different from the mechanical picture. In this sense, in our protein model the mechanical and thermal descriptions provide incompatible views of the folding/unfolding landscape of the system, and the estimated quantities to zero force result hard to interpret.

## I. INTRODUCTION

Force spectroscopy techniques have provided a wealth of high resolution data, which resulted in a great step forward towards the understanding of how proteins fold^1,2^. Following the classical meaning of spectroscopy, a biomolecule is perturbed by an external mechanical bias that induces the unfolding transition, and from which information from the biomolecule folding landscape can be inferred. For example, in a force extension protocol, a probe such as an AFM cantilever is retracted at constant velocity, applying an increasing force that eventually unfolds the protein^3,4^. From the relationship between the unfolding force and the pulling velocity, we can learn about the mechanical stability, or the unfolding pathway of the subject molecule, including how it is modulated by physiological factors, such as disulfide bonds^5^, or chaperones^6,7^. In addition, the implementation of force-clamp techniques allowed to apply constant forces to individual proteins and record the unfolding time of the perturbed system^8^, but also to carry out equilibrium folding/unfolding experiments and explore in great detail the folding pathways^9,10^, or rare events which might appear over long timescales^11–13^.

In this context, many theoretical models have been developed to provide tools to analyze the experimental data and obtain physical quantities about the system. Roughly, we can divide such theoretical efforts into those devoted to the kinetics, and those to the equilibrium properties. Bell-Evans model provided a first phenomenological framework which allowed to recover the distance to the transition state and the unfolding rate at zero force from force-extension and constant force experiments^14,15^. Later refinements of this theory allowed to recover also the height of the free energy barrier at zero force^16–18^. On the other hand, the famed Jarzynski equality provides the estimation of equilibrium free energy quantities^19^, and even allows to reconstruct the full free energy profile from such non-equilibrium work measurements^20^. Importantly, in both cases we obtain the estimation of free energy magnitudes extrapolated to zero force, and thus it appears tempting to compare them directly with those obtained from biochemical experiments, such as thermal of chemical denaturation^3,21,22^.

However, an evident question arises in this context; how meaningful is such comparison? When folding and unfolding mechanically, the pulling force imposes a very specific reaction coordinate, which is the pulling vector. As protein folding is known to be a highly complex and multidimensional process, it is unlikely that the distance between the N and C-terminal of a protein provides a relevant order parameter to describe protein folding in the absence of force. Even more, while at very high forces a protein will unfold along the pulling directions, this is not clear to be the case at low forces, where orthogonal degrees of freedom might be of relevance, compromising the use of the aforementioned theories to explain the data.

Here, we explore these questions by analyzing in high detail the unfolding landscape of the BPN_46_ model protein when denatured by temperature and under a mechanical bias. The BPN_46_ protein is a highly studied non-Go protein model, due to its rich and complex landscape. It folds spontaneously into a *β* barrel-like structure, that shows frustration in the specific arrangement of its *β*-sheets, having also intermediate conformations and a non-trivial folding mechanism when subject to force^23–27^. Inspired by typical force spectroscopy protocols, we first unfold the protein mechanically using non-equilibrium force extension and constant force modes. These protocols measure the unfolding forces and the unfolding rates as a function of the pulling velocity and the pulling force, respectively. By analyzing this dependence, the extrapolated to zero force distance to the transition state and the height of the free energy barrier can be estimated, assuming a simple one-dimensional pathway^16–18^. Furthermore, the extended Jarzynski equality can be used to reconstruct the zero-force free energy profile by sampling the individual force extension trajectories^20^. Our data shows that the protein unfolds by surmounting two distinct free energy barriers, first reaching a mechanical intermediate conformation, previously identified^26,27^. Using the lowest pulling velocity to reconstruct the free energy profile, we obtain a similar unfolding picture, but with lower free energy barriers, likely to the multiple unfolding pathways that compete at very low forces, as previously reported^27^.

In order to explore the folding landscape in the absence of force, we carry out equilibrium simulations in the vicinity of the critical temperature. By using Markov state models and transition state theory, we obtain a detailed description of the conformation landscape of the system, and the unfolding pathway. While we can relate structurally those states excited mechanically and thermally, their role in the protein dynamics seems to be completely different. The thermal unfolding mechanism can be fairly well described as a one-directional pathway with two well-defined intermediate states, different from those observed in the mechanical picture. While our conclusions are specific to the BPN_46_ protein model, which has some non-standard features for a simple protein folding model, our data exemplifies some of the problems that might arise when directly comparing data obtained from different denaturants. Protein folding in presence and absence of force are very different processes; hence, it is unclear how extrapolations to zero force inform about a folding transition in the absence of force.

## II. MODEL AND SIMULATION METHODS

The BPN_46_ model is a coarsed-grained non-Go protein model^23–26,57^. It has a 46-residue sequence where each residue is represented as a “colored” particle, being either hydrophobic, hydrophilic, or neutral. The Hamiltonian of the model is defined by four potential terms: a stiff nearest-neighbor harmonic potential, a three-body bending interaction, a four-body dihedral interaction and a sequence-dependent Lennard-Jones potential. This latter term contains the sequence dependency, since hydrophobic residues attract to each other while every other pair see a short-range repulsive potential.

The BPN_46_ folds spontaneously into a stable native structure as a *β*-barrel constituted by four *β*-strands and three neutral turns (see Fig. 1A), mimicking the fold of other well-known domains. However, potential energy analysis of the protein conformational landscape have revealed a multiplicity of structurally similar ground states, separated by large barriers^25^. The structure is held together by the interaction between strands *β*_1_ and *β*_3_, formed by just hydrophobic residues. Figure 1B shows the map of native contacts. The hydrophobic strands *β*_1_ and *β*_3_ run in a parallel disposition, while strands *β*_1_-*β*_2_, *β*_2_-*β*_3_ and *β*_3_-*β*_4_ run antiparallel between them.

**FIG. 1.**
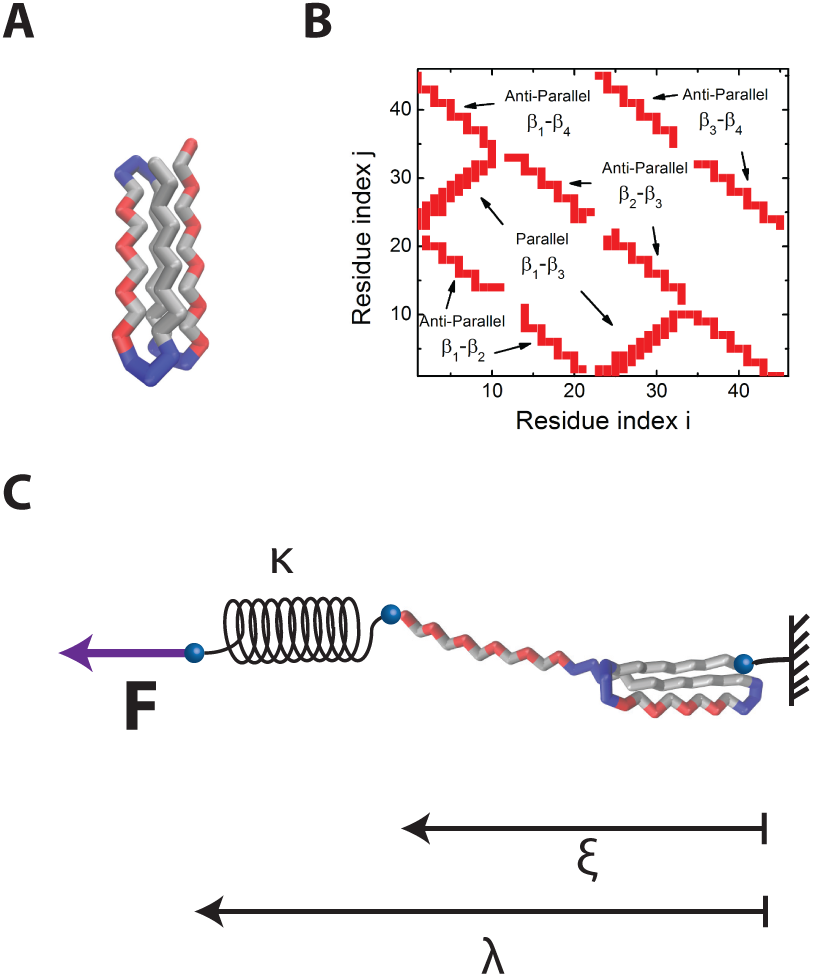
The BPN_46_ model protein. (A) Representation of the native structure of the BPN46 model protein. Hydrophobic residues (B) are shown in grey, neutral (N) in blue and hydrophilic (P) in red. (B) Map of native contacts of the BPN_46_ model protein. It has a *β* barrel structure arranged in 3 antiparallel and one parallel *β* sheets. (C) Configuration for the pulling simulations. The protein is fixed from one end while from the other end is attached to a linear spring of constant *κ*. Two different pulling protocols are carried out. In the force extension one, the *λ* coordinate is linearly increased *λ* = *vt*, where *v* is the pulling velocity. In the constant tween the ends of the protein. In contrast with *λ* which is a control parameter, *ξ* represents the end to end distance of the protein and is a stochastic magnitude.

The behavior of this model protein as a function of temperature and force was reported before^24,26–28^. It exhibits a unfolding transition, with a well-defined peak on its heat capacity at a temperature *T*_*c*_ = 0.9*kT*.

In all simulations, we generate stochastic trajectories by integrating numerically the Langevin equation of motion for the 46 residues,

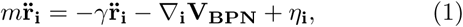

where *m* is the unitary mass of each residue, *γ* the friction coefficient, *V*_*BPN*_ the potential defined for the BPN_46_ model (see App. A), and *η*_*i*_ a gaussian white noise term of zero average, which fulfills the fluctuation-dissipation theorem ⟨ *η*_*i*_*η*_*j*_⟩ = 2*kTγδ*(*t* − *t* ′)*δ*_*ij*_.

In the mechanical simulations, we carry out non-equilibrium unfolding by fixing the N-terminal of the protein and attaching a linear spring of constant *κ* to the C-terminal, used to apply force (see Fig. 1C). Force extension trajectories are generated by displacing the spring at a constant velocity *v*, so that the control parameter *λ* increases linearly *λ* = *vt*. Constant force trajectories are generated by suddenly displacing the spring by an amount Δ*x* so that it generates a constant force *F* = *κ*Δ*x*. In all mechanical simulations, we work at a temperature of 0.55*Tc*. When working at this temperature the protein unfolds mechanically at a force of *F*_*U*_ ≈ 20 pN. Here, we will study nonequilibrium unfolding trajectories at forces above the unfolding force. A conceptually different case is that of the folding-unfolding equilibrium dynamics at a force below the unfolding force, studied in detail in^27^ (there, *T* = 0.55*T*_*C*_ and *F* = 0.8*F*_*U*_).

Equilibrium thermal folding-unfolding simulations are carried out at the vicinity of the transition temperature (1.1 *T*_*c*_), to ensure the system visits the maximum number of configurations.

## III. MECHANICAL UNFOLDING

### A. Analysis Methods

#### a. Force Extension

In force extension trajectories, the experimental output is the force as a function of the pulling coordinate *λ* = *ξ* + *F/κ*. Molecular transitions are identified as rupture peaks characterized by the rupture force, which increases with the loading rate *r*_*f*_ = *vκ*. Bells-Evans phenomenological model predicts a logarithmic dependence of the average rupture force with the loading rate ⟨*f*\* ⟩ ∼ log(*r*_*f*_)^14,15^. Extensions of this model predict ⟨ *f \**⟩ ∼ log(*r*_*f*_) ^ν^, wher *ν* is an exponent that depends on the potential shape chosen to model the irreversible rupture process: *ν* = 2*/*3 for a linear-cubic, *ν* = 1*/*2 for a parabolic-cusp and *ν* = 1 for a linear potential, which recovers Bell-Evans model^18^. Generally, it can be argued that the linear-cubic landscape is more appropriate, as close to the transition any analytic potential tilted by a pulling force can be approximated by a linear-cubic potential. By fitting the average rupture force ⟨ *f* * ⟩ to the loading rate, the free energy barrier height Δ*G*^†^ and position *x*^†^, and the intrinsic rate *k*_0_ can be obtained as:

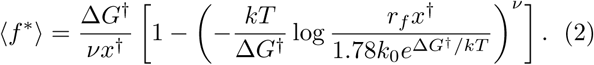

#### b. Constant Force

In constant force trajectories, the experimental output is the end-to-end distance *ξ* as a function of the simulation time. Molecular transitions are identified as discrete increases in the extension, characterized by the rate of rupture *k*, which increases with the pulling force. In a similar way to what described above, Bell-Evans model predicts an exponential dependence of the rate with the pulling force. Extensions of the model predict a more complex behavior, and fitting *k*(*F*) allows recovering the free energy barrier height 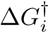 and position 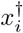 and the intrinsic rate *k*_0_:

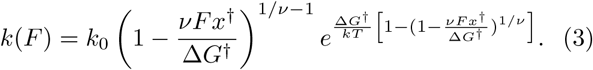

The rate of rupture can be estimated by averaging all rupture traces and fitting to a single exponential or multiple exponentials, if more than a kinetic process is involved.

#### c. Reconstruction of the free energy profile

From non-equilibrium pulling experiments, the free energy profile can be reconstructed by using the extended Jarzynski expression^20^:

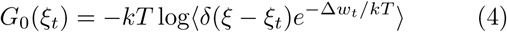

where *G*_0_(*ξ*) is the free energy profile at zero force along the pulling coordinate *ξ*, and Δ*w*_*t*_ the difference between the external work done on the system, and the biasing potential, 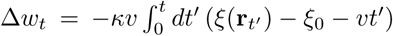. In this sense, we take time slices and average over different trajectories the work necessary to take the system there, and estimate the free energy at that time slice.

### B. Results

#### a. Force extension

We carry out force extension simulations at pulling velocities between 0.002 and 0.2 *nm/τ* ^29^, recording the dependence of the force with the pulling coordinate *λ*. Figure 2A shows typical force extension trajectories for *v* = 0.004 nm*/τ*. Light grey lines show three individual realizations, while the black curve represents the average over a total of 100 trajectories. The protein extends through two different transitions, as every trajectory shows two peaks. We identify in the average a first broad low force transition which leads to *λ* ∼11 nm, and a second high force one leading a state with *λ* ∼ 22 nm. We can relate the first state to the half-stretched conformation (HS), while the second to the fully extended conformation, following the notation previously stablished^27^.

**FIG. 2.**
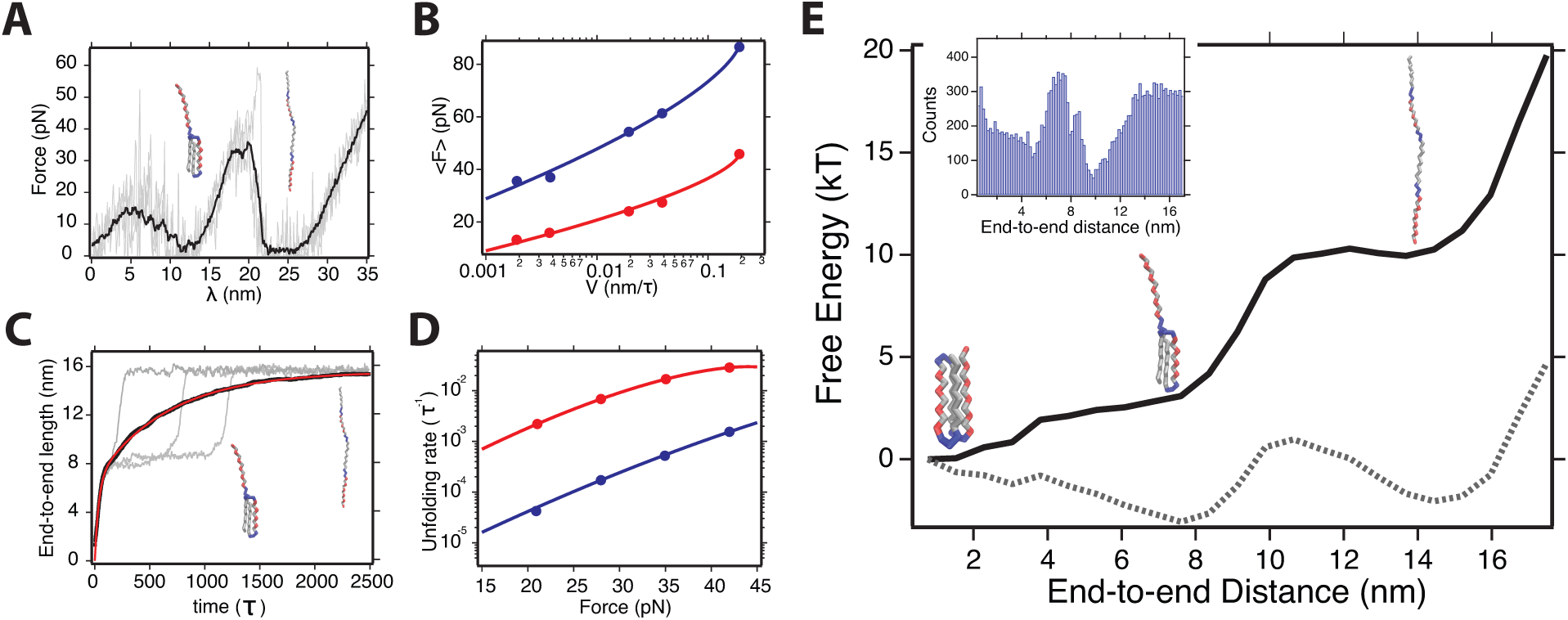
Non equilibrium mechanical unfolding simulation results (in all cases *T* = 0.55*T*_*c*_). (A) Typical force extension trajectories at *v* = 0.004 nm*/τ*. Grey lines show individual realizations, and black line the average unfolding trajectory over 100 realizations. The mechanical unfolding in the force extension protocol occurs through two well distinguished transitions, a first low force one at *λ* ∼ 5 nm and a second high force one at *λ* ∼ 17 nm. (B) Average unfolding force as a function of the pulling velocity for the first transition (red) and second transition (blue). Data points are fit to the Eq. 2 with *ν* = 2*/*3, obtaining 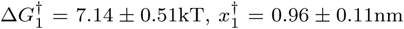 and 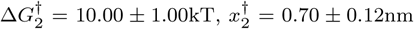. (C) Constant force trajectories at 42 pN. Grey lines correspond to individual realizations, and black line is the average over 100 realizations. Analogously to force extension experiments, unfolding occurs through two subsequent transitions, one yielding to an extension of ~ 8nm and the second to ~ 16nm. Red curve is a fit to a two-exponential model, yielding the microscopic rates for the two kinetic transitions. (D) Unfolding rates as a function of the pulling force for the first (red) and second (blue) transitions. Solid lines are fits to Eq. 3 with *ν* = 2*/*3, obtaining 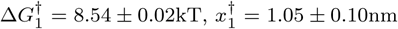 and 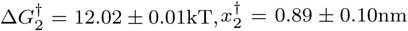. (E) Free energy profile along the end-to-end distance reconstructed from force-extension trajectories at *v* = 0.002 nm/*τ*, using the extended Jarzynski equality. The profile yields a similar picture, where the unfolding process takes place through a mechanical intermediate with and end-to-end extension of ∼7 nm—HS conformation. However, the free-energy barrier estimated to reach the intermediate is lower than that using the complete pulling rate range, likely since, due to the low forces involved, multiple pathways are involved, as previously reported. (Dotted line) Tilted landscape under a force of 18 —pN. (Inset) Residence time histogram along the end-to-end distance, from which the profile is built.

Figure 2B shows the rupture forces fitted to Eq. 2, using *ν* = 2*/*3 where red corresponds to the first event and blue to the second one. Thus, according to this description, there is a first barrier characterized by 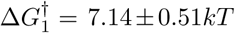 and 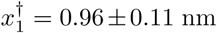, and a second one with 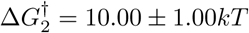 and 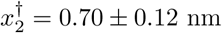.

#### b. Constant force

We carry out constant force simulations at forces ranging from 20 to 42 pN. Figure 2C shows constant force trajectories where the end-to-end distance *ξ* is plot against the time for a pulling force of 42 pN. Light grey traces correspond to three individual realizations, while the black trace is the average trajectory obtained from averaging 100 realizations. Similarly to the force extension protocol, unfolding occurs sequentially through a mechanical intermediate, with an end-to-end distance of, *ξ* ∼8 nm, in accord to the extension of two *β* strands. As unfolding occurs through two kinetic process, we represent the average extension as a double exponential ⟨*ξ* ⟩(*t*) = *ξ*_1_(1 − exp(− *k*_1_*t*)) + *ξ*_2_(1 − exp(−*k*_2_*t*)), where *ξ*_1_ is the extension of the mechanical intermediate and *ξ*_2_ the length increment to the unfolded states, being *k*_1_ and *k*_2_ the kinetic rates of the two pro-ceses, respectively.

Unfolding rates obtained at different pulling forces are fitted to Eq. 3, with *ν* = 2*/*3, where red corresponds to the first barrier and blue to the second one. We obtain 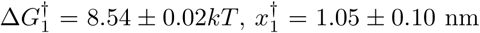 for the first barrier 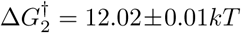, and 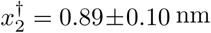 for the second one. The values are in agreement with those obtained for the force extension protocol.

#### c. Reconstruction of the free energy profile

Finally, we use the force-extensions trajectories to reconstruct the free energy profile along *ξ* by using the extended Jarzynski equation (Eq. 4)^20^. Here, taking *ξ* = 0 as the reference distance, the free energy at different values of *ξ* is estimated from the exponential average of the non-equilibrium work required to reach that distance. Jarzynski equality has well-known convergence problems, in particular when the data used is very far from equilibrium—the number of trajectories needed for convergence increase with the exponential of the dissipated work. Hence, we use the slowest pulling speed (*v* = 0.002 nm/*τ*) for the reconstruction. This landscape is an estimation of the free energy for the unperturbed system, and thus should be directly comparable to the free energy barriers calculated through Eqs. 2 and 3.

Figure 2E shows the reconstructed profile (solid line). The unfolding mechanism suggested here agrees with that depicted by the force-extension and constant force simulations, since the unfolded state is reached through a mechanical intermediate—HS configuration. These three energy minima are more evident when tilting the landscape with a pulling force (dotted line). The extensions of the three involved states match the plateaus at constant force (Fig. 2C), with the native state with a ∼0 nm end-to-end length, the HS with ∼7 nm, and the extended state at ∼14 nm. The unfolded state is reached from the HS state after surmounting a ∼8 kT, similar to that estimated in the fits of Figs. 2 B and D. By contrast, the native and HS state appear separated by an almost negligible barrier, while the fits to Eq. 2 yielded a barrier of ∼7 kT. This discrepancy arises likely due to the very low forces at which the transitions occur at this low pulling velocity. As reported at low constant forces^27^, the conformational landscape of the BPN_46_ protein under force is complex, and rich dynamics appear between the native and HS state, with several other metastable states separated by very low barriers. Hence, it is likely that at very low pulling speeds, these quasi-equilibrium transitions take place and average out the free energy profile in the Jarzynski reconstruction, yielding to a lower free energy barrier. Indeed, while the average force-extension trajectory at *v* = 0.002 nm/*τ* shows two clear peaks at forces of∼15 and ∼35 pN, inspection of individual trajectories reveal complex patterns of unfolding peaks between 0 and 8 nm, suggesting complex dynamics, compatible with those that appear at a constant low force^27^.

## IV. THERMAL UNFOLDING

### A. Thermal unfolding simulation details

We carry out five simulations with a duration of *t* = 10^9^*τ* each, close to the critical temperature *T* = 1.1*T*_*c*_. Each simulation is previously thermalized during *t* = 10^4^*τ*, to randomize the initial conditions of each trajectory. The simulation time is sufficient for the system to adequately explore its conformational landscape in equilibrium, given the high temperature, which leads to very fast dynamics between all visited states. Equilibrium is later checked on the Markov network, showing that detailed balance holds.

### B. Analysis Methods: Dimension reduction, Markov state model and transition-path theory

In order to analyze the equilibrium thermal folding-unfolding process, we represent the configurational space of the BPN_46_ protein at *T*_*c*_ as a Markov state network^30^. In this representation, the free energy landscape is shown as a complex network, where the nodes correspond to macrostates (free energy basins) obtained by some clustering method, while the edges—weighted and directed— relate to the kinetic transitions between such states. In a nutshell, building a Markov state network starts by a geometrical discretization of the configurational landscape to build a first microstate network, typically with thousands of nodes, and hence, hard to interpret. Next, this network is coarse-grained by some lumping algorithm, that clusters those states kinetically related, to end up with a smaller, more significant representation of the system’s energy landscape. In our case, we use the SSD algorithm as a clustering method^27,31,32^.

The full configurational landscape can have typically several hundreds of dimensions, and a direct discretization of it is a futile effort; hence, it is useful to reduce first the dimensionality of the system to a few, meaningful coordinates. These coordinates should yet be able to identify the large and slow conformational transitions, which would allow to rule out the abundant and meaningless fast fluctuations. Principal Component Analysis (PCA) is often used as a dimension reduction method^27,31–33^, since it identifies the coordinates that contain the largest structural fluctuations about the average conformation. However, it has been recently demonstrated that Time-structure Independent Component Analysis (TICA) is the optimal method for identifying the ‘slow” order parameters, since it takes into account, not only the spatial variation between structures, but also the timescales over which they occur^34,35^. In our previous work^27^, PCA was sufficient to separate the main macrostates occupied by the system along its dynamics. However, those simulations were carried out at low temperature, and under force; hence, the dynamics where intrinsically slow, given the large free energy barriers that separated the free energy basins of the system. Here, we carry out simulations close to the critical temperature in order to populate the unfolded state without the need of an external bias. Therefore, the dynamics and transitions that define the system dynamics are much abundant and fast, and PCA is a method likely to fail in separating the main conformational changes of the system.

Figure 3 compares the projection of the simulated dynamics along the first two PCs (A), and the first two TICA components (B). These two components capture the three main conformational states of the system, the native (N), the half-extended (HE) and the unfolded state (U). However, it is very evident by comparing both projections, that TICA does a much better job in defining the energy basins and free energy barriers that separate them. The N and HE states are separated by a ∼2.5 kT barrier in the PCA projection, while this barrier is of ∼4 kT in the case of TICA. The unfolded conformation— with a low population—only appears as a spatially spread state (given its large entropy) in the TICA projection, while its representation in the PCA projection is much dimmer. Finally, along the second TICA coordinate, the HE shows multiple states associated to similar conformations separated by small energy barriers, that are not separated with PCA. Hence, this demonstrates the need to use TICA to find meaningful order parameters to build the Markov network.

**FIG. 3.**
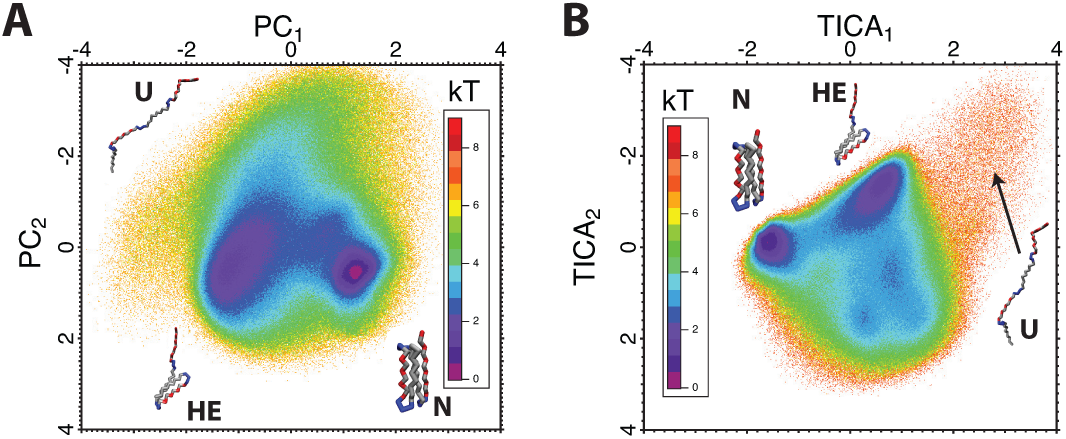
Comparison between PCA and TICA as dimension-reduction methods. (A) Two-dimensional landscape projected along the first two principal components. (B) Two-dimensional landscape projected along the first two TICA components. Compared to PCA, TICA allows to obtain better defined states that facilitate the discretization and clustering of the trajectories, and the calculation of the Markov State network.

We use the first three TICA components as our configurational space, and discretize them into 30 bins, obtaining a microstate network with 6657 nodes connected through 228682 links. We then lump the microstates onto kinetically significant macrostates by applying the SSD algorithm. We obtain a first ‘raw” macrostate network with 45 basins of attraction of macrostates. We refine such network, eliminating basins with an occupation *π*_*i*_ < 10^−4^, to avoid extremely rare or pathological states. The final macrostate network is made up of 21 nodes, and we take it as the representation of the free energy landscape of the system.

From the equilibrium Markov state network, we can calculate precisely the unfolding pathways by applying transition-path theory^36–38^. Briefly, we start defining the set of native conformations (N) and the set of unfolded conformation (U). Next, we will rank all other states, as intermediates I between N and U, depending on how close they are to the unfolded conformation in terms of unfolding pathways. To this aim, we calculate the committor probability (or unfolding probability) 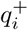 of each state (being 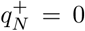 and 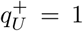 by definition) as the solution for the system of equations:

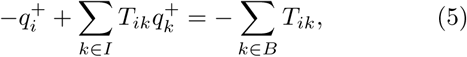

where *T*_*ik*_ is the rate matrix of the Markov network. Intuitively, 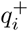 represents the probability that, state *i* reaches the unfolded conformation U before reaching back the native state N. Then, we compute the unidirectional flux to the unfolded state as 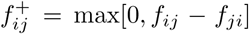, where 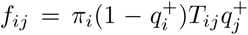. 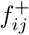 defines a network of fluxes from the native N to the unfolded U configurations, from which we extract the individual unfolding pathways.

#### a. One dimensional free energy landscapes

Figure 4 shows a fragment of the simulated trajectories projected along three different order parameters, the first and second TICA components (Figs. 4A and C), and the fraction of native contacts (Fig. 4E). These fragments represent only 1% of the total simulated time, which demonstrates that the system explores its landscape in a much faster timescale than the simulated time window. Right panels are the one-dimensional landscape as obtained from each coordinate. The first TICA component separates the transitions between the three major states, the native (N), half-extended (HE), and unfolded (U) states. Most of the dynamics correspond to very fast transitions between the (N and (HE) states, with occasional visits to the unfolded state. The (N) state has a narrow free energy basin (Fig. 4B), while the (HE) state appears as shallower basin, likely due to the presence of multiple conformations, that will be resolved in higher order TICA components. This suggests that, even though the native state of the system is frustrated, such frustration is removed at the simulation conditions, likely due to the high temperature that sheds the free energy barriers between the multiple ground states. The second TICA coordinate maintains the transitions to the unfolded state, while it shows a much richer conformational landscape, resolving more subtle structures around the (HE) state. Finally, the fraction of native contacts (Q), is useful to identify the states populated by the dynamics, since it indicates intuitively the degree of nativeness of the states. However, it does not provide a good reaction coordinate to represent the dynamics, since most minima are shallow basins, separated by small barriers, suggesting that multiple states might be averaged out when projecting onto this coordinate.

**FIG. 4.**
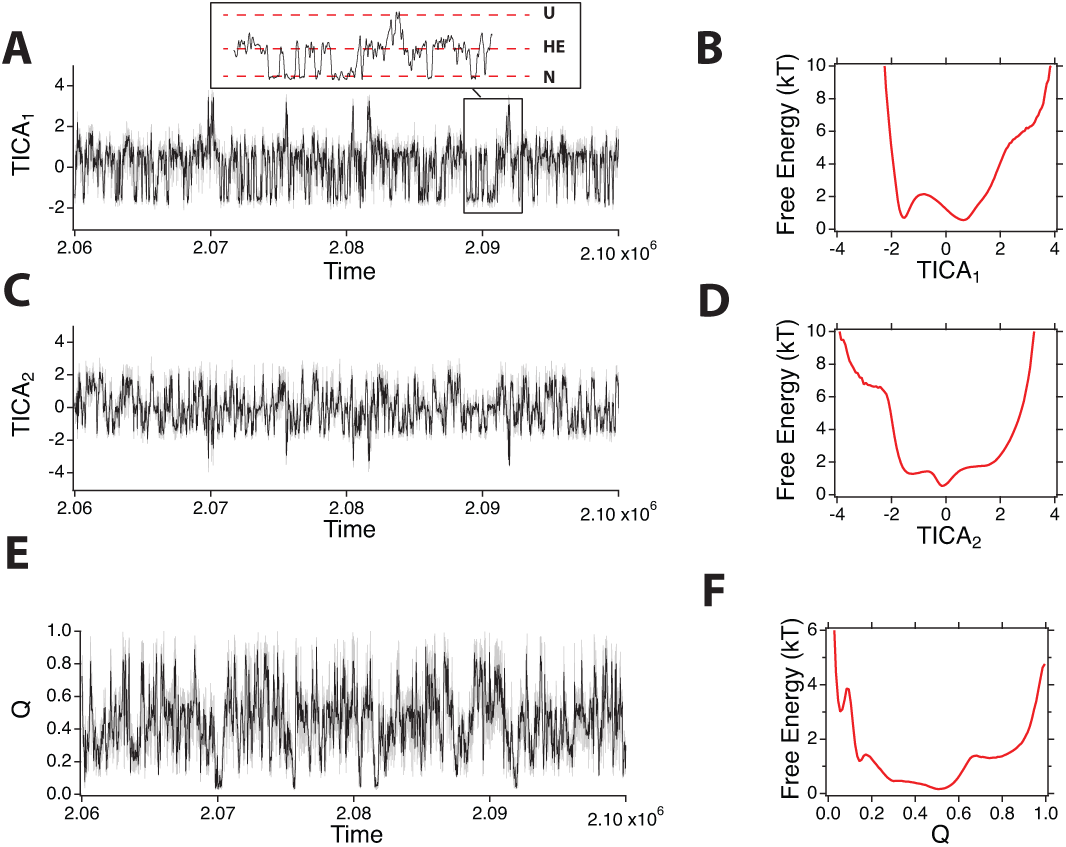
Fragment of the simulated trajectories along different order parameters, with their projected one-dimensional landsacpe. (A) Trajectory along the first TICA coordinate. (Inset) detail where the three main states are identified. (B) One-dimensional landscape along the first TICA coordinate. (C) Trajectory along the second TICA coordinate. (D) One-dimensional landscape along the second TICA coordinate. (E) Trajectory along the fraction of native contacts. (F) One-dimensional landscape along the fraction of native contacts.

#### b. The equilibrium Markov state network

Figure 5A shows the macrostate network, calculated as explained above. The size of the nodes is proportional to their population with a cutoff of *π*_*i*_ < 10^−2^, below which beads have the same size, for visualization reasons. Links with *T*_*ij*_ *>* 10^−5^*τ* ^−1^ are represented as arrows connecting states. We calculate the fraction of native contacts *Q* of each node (see Table 1), which allows us to cluster them into six major regions, represented in different colors: The native state (N) (*Q* ≈ 0.8; blue); the half-extended configuration (HE) (*Q* ≈ 0.5; red); the collapsed ensemble (C), (*Q* ≈ 0.3; green); the first intermediate state (I_1_) (*Q* ≈ 0.2; yellow) the unstructured ensemble (UE) (*Q* ≈ 0.2; purple), the second intermediate state (I_2_) (*Q* ≈ 0.1; magenta), and the unfolded ensemble (U) (*Q* ∼ 0; black).

**TABLE I.** Magnitudes characterizing the states detected in the equilibrium network. We characterize the states with their occupation *πi*, fraction of native contacts *Q*, unfolding probability, and mean escape time *t*_*e*_ (in *τ* dimensions).

| State | $\pi_i$ | $Q$ | $p_{\text{unfold}}$ | $t_e$ ($\tau \times 10^3$) |
| --- | --- | --- | --- | --- |
| N | 0.291 | 0.82 | 0.00 | 14.30 |
| HE | 0.372 | 0.51 | 0.00 | 13.70 |
| C | 0.264 | 0.33 | 0.00 | 6.25 |
| $I_1$ | 0.068 | 0.23 | 0.14 | 3.50 |
| UE | 0.002 | 0.19 | 0.27 | 1.53 |
| $I_2$ | 0.036 | 0.14 | 0.46 | 3.92 |
| U | 0.002 | 0.06 | 0.95 | 1.80 |

**FIG. 5.**
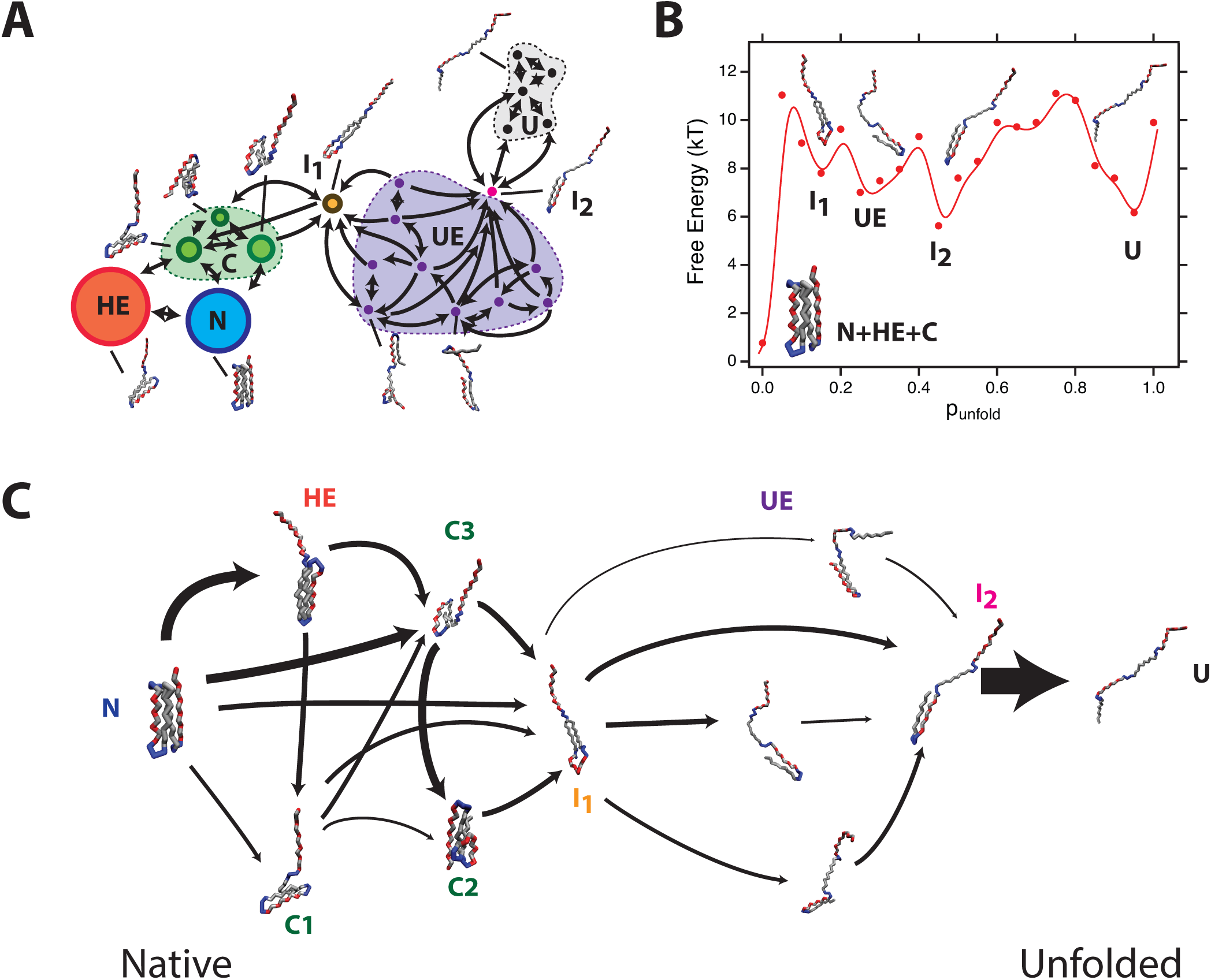
Markov state model of the BPN_46_ model protein close to the critical temperature. (A) Conformational macrostate network at *T* = 1.1*T*_*c*_. Each macrostate is represented as a bead which size is proportional to its population (although states with population *π <* 10^−2^ are represented as beads of same size). Arrows represent observed transitions between states. Most representative structures are shown next to the associated state. The free energy landscape can be clustered in six major conformational regions, represented by the colors in the beads. (B) Free energy profile along the *p*_unfold_ reaction coordinate. Data points are the clustered histogram from projecting the Markov network onto the reaction coordinate, and the solid line is an interpolation to depict a smooth landscape. (C) Folding pathways as obtained by applying TPT to the macrostate network. Folding/unfolding occurs in a very one-dimensional fashion, being bottleneck states of the unfolding pathway.

When comparing with the unfolding mechanism described for the non-equilibrium pulling simulations, besides the native and unfolded state, we find the HE configuration, which resembles the mechanical intermediate described in the landscape of Fig. 2E. Both the HE (thermal) and HS (mechanical) states maintain the core structure and show strand *β*4 extended. In the thermal network, the HE conformation has *π*_*HE*_ = 0.53, and is heavily connected to the native conformation, with fast transitions between them *T*_*HE,N*_ ≈ 3 × 10^−5^*τ* ^−1^. This is similar to the almost barrierless landscape we encountered in the mechanical description in Fig. 2. Additionally, we identify two states that play a central role in the folding/unfolding dynamics; state I_1_—which maintains the hydrophobic interactions present in the native interactions between *β* strands 1 and 3, providing 30% of the native contacts—and the state I_2_—which maintains little structural similarity with the native state, since the hairpin formed by the weak interactions between the hydrophobic residues in *β* strands 1 and 2 is the only motif that survives from the native conformation. These two states are bottlenecks in the pathways connecting the native and unfolded states, and removal of any of them will disconnect both ensembles.

#### c. Free energy profile along the commitor probability

The committor probability 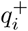, or unfolding probability *p*_unfold_, is often argued to be the appropriate reaction coordinate for describing (un)folding transitions^37^. In this sense, we can project the simulated trajectories onto *p*_unfold_, and calculate a free energy profile along it.

Figure 5B shows the free energy profile along *p*_unfold_, where the points correspond to actual free energies estimated by binning *p*_unfold_ with a bin size of 0.05, and the solid line is an interpolation to visualize a smooth landscape. All states belonging to the native (N), half-extended (HE) and collapsed (C) ensembles appear lumped onto the same free energy minimum, with *p*_unfold_ ∼ 0. This indicates that the majority of the system dynamics involves internal transitions among these three ensembles, that do not lead to unfolding. The intermediate state (I_1_) is the first relevant minimum, with *p*_unfold_ ∼ 0.15. This procedure allows us to identify the transition state as that with 50 % probability of unfolding, which corresponds to the state labeled as I_2_ in the landscape network.

#### d. Unfolding pathway

By applying TST, we can determine the unfolding pathways by converting the Markov state network shown in Fig. 5A into a one-directional flux network that connects the native state (N) with the unfolded ensemble (U). Figure 5C shows the unfolding network, depicted from left to right in terms of increasing *q*^+^, where arrows connecting states have a thickness which is proportional to their flux 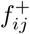, being only those connections with 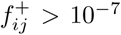 represented for visualization reasons.

As suggested by the landscape network, thermal unfolding occurs in a one-dimensional way, where states I_1_ and I_2_ are bottleneck nodes, that concentrate the unfolding flux. Most of the dynamics between states N, HE and C correspond to conformational changes that do not drive unfolding. However, we can identify states C2 and C3 (belonging to the collapsed ensemble) as precursors of the intermediate I_1_, that eventually leads to state *I*_2_, and the unfolded state. In this sense, the flux network suggests that the folding/unfolding equilibrium transitions of the protein follow a one-dimensional dynamics, where *p*_unfold_ works indeed as a good reaction coordinate. This unfolding dynamics, although one dimensional as that described in Fig. 2 when applying large forces, follows a very different pathway from the nonequilibrium mechanical unfolding. The half-stretched conformation–to which HE can be ascribed as the non-tensioned homologous– played a very clear role as a mechanical intermediate, as shown in the free-energy profile reconstruction. However, the one-dimensional landscape here has a very different structure, and the state HE, although kinetically relevant, does not play a significant role in the unfolding pathway. Interestingly, this simple pathway contrasts greatly to the dynamics exhibited by the protein when a small mechanical force is applied. Mechanical forces seemed to excite conformations that do not appear in the absence of force, and thus created a much more complex unfolding dynamics, that could not be described through a one-dimensional landscape^27^.

## V. DISCUSSION

The results presented in this work, together with those previously published^27^, present a threefold vision of the unfolding mechanism of the BPN_46_ model protein; while we previously described the equilibrium landscape at low force, here we present the landscape extrapolated to zero force from non-equilibrium pulling simulations—analogous to what often done in force spectroscopy experiments—and an equilibrium landscape obtained from thermal simulations. When comparing these three landscapes, some similarities appear between the unfolding processes, and, more notably, irreconcilably differences.

The BPN_46_ is a simple protein model, that, however, exhibits a complex and rich conformational landscape, including a frustrated ground state, intermediates, or a multidimensional folding mechanism^26,27,39,40^. Therefore, our results must be understood as an exemplification of the dichotomy that might arise when comparing folding mechanisms through different denaturants, such as force and temperature. Interestingly, in every of the unfolding protocols we have used, we recover a similar metastable state, which we dubbed the HS configuration in the mechanical simulations, and HE configuration in the thermal landscape. This state maintains the hydrophobic core between *β*-strands1-3, with the *β*-strand4 dislodge from the structure. The prevalence of this conformation in every landscape illustrates a similarity between the landscapes recovered through different techniques. However, the role of this state in the folding/unfolding mechanisms is very different, depending on the protocol we use. In non-equilibrium unfolding simulations, the HS conformation appears as a clear mechanical intermediate, and the pulling trajectories reflect the need to surmount two free energy barriers to reach the stretched state (Fig. 2). These barriers can be estimated using the Dudko-Hummer-Szabo theory^18^, resulting in a ∼7-8 kT barrier separating the native and HS conformation. However, in the absence of force, this analogous state appears as a relevant metastable with little role in the unfolding mechanism, similar to what found in equilibrium simulations at very low forces^27^. This indicates that the mechanical regime used to trigger unfolding can also alter the pathway the protein explores, preventing the use of extrapolations to zero-force. Indeed, the reconstruction of the one-dimensional profile using the extended Jarzynski equality reveals a small first barrier, likely due to the competence of several transitions that occur at very low force.

In this sense, the role of mechanical forces in driving protein unfolding must be extrapolated with caution, not just when establishing parallelisms or comparison with unfolding driven by other denaturants (either thermal or chemical), but also when comparing mechanical unfolding under different protocols. Here, we have shown how, surprisingly, the thermal unfolding mechanism of the BPN_46_ protein is rather simple; while it exhibits rich dynamics between different metastable states such as the HE or the C ensemble, the protein unfolds in a one dimensional way, with two well defined intermediate states (I_1_ and I_2_). However, when low forces are applied such that we allow equilibrium transitions between the folded and unfolded states—similar to what it is possible to do with force-clamp spectroscopy techniques—the unfolding mechanism is rather complicated. Contrary to what could be intuitively predicted—since the force should impose a reaction coordinate and thus the unfolding mechanism could be easily described by this geometrical coordinate—the unfolding mechanism gets more complicated, and roughly two major unfolding routes can be identified^27^. When the magnitude of the force is increased so that the equilibrium is shifted to the unfolded conformation, the intuitive behavior arises, and the protein unfolds through the end-to-end reaction coordinate. This implies that the effect of force on the landscape of this protein cannot be modeled over the whole range as a simple linear perturbation with a −*fx* term, as commonly done.

Since the advent of force spectroscopy techniques, free energy quantities extracted from pulling experiments have been usually contrasted with those extracted from bulk experiments, such as chemical or thermal denaturation^3,22,41–43^. This comparison implicitly assumes a simple two-state one-dimensional folding landscape, which, while can be reasonable for some systems over a certain scale, should not be generalized. Force is a unique denaturant, since it imposes a topological constraint on the tethered molecule, and defines a very specific reaction coordinate, which might not be relevant in the absence of force. Indeed, mechanical pulling experiments and simulations have been shown to reproduce very different dynamics compared to those exhibited in the absence of force^21,44–48^. For instance, downhill proteins are relevant for folding in solution over a marginal free energy barrier, contradicting the classic two-state picture^49–51^. However, recent force spectroscopy experiments on the gpW downhill protein showed that, under force, this proteins transitions between the folded and unfolded state with two-state dynamics over a barrier of about 2.5 kT^52^. In this sense, recent instrumental development have increased the resolution at which we can interrogate a folding protein, revealing a much more complex scenario than what typically assumed. Protein L, a very classic two-state folder, folds through a ephemeral molten globule-like state that lasts few milliseconds, and therefore is only captured through over very short force quenches^9^. Additionally, the new access to very long timescales, allowed exploring unfolding at low forces, and showing non-monotonic dependences of the unfolding rates with force, which can be interpreted as a multipathway unfolding mechanism^53,54^. Therefore, while our studies here deal with a specific protein model with very particular features, our data exemplifies the dichotomy that can arise, not only when unfolding mechanically and thermally a protein, but also when understanding this mechanism over different force ranges.

In summary, our works provides a practical case of a simple protein that, however, exhibits complex unfolding dynamics that depend strongly on the protocol used to unfold it. Non-equilibrium mechanical protocols as those typically used in single molecule pulling experiments reveal one-dimensional unfolding through a mechanical intermediate, that, however, does not play any role in the unfolding pathway, neither at low or in the absence of force.

## ACKNOWLEDGMENTS

This work was supported by Spanish Ministerio de Economía, Industria y Competitividad (MINECO) Projects No. FIS2014-55867-P and FIS2017-87519-P, co-financed by the Fondo Europeo de Desarrollo Regional (FEDER), and by the Gobierno de Aragón, Grant to the FENOL group, E36 17R.

## Appendix A: Model parameters and simulation procedures

We use the same model protein employed in^27^. The interaction between residues account for four terms, an stiff harmonic nearest neighbor interaction, a three-body bending interaction, a dihedral four-body interaction, and a sequence-dependent Lennard Jones potential, this is:

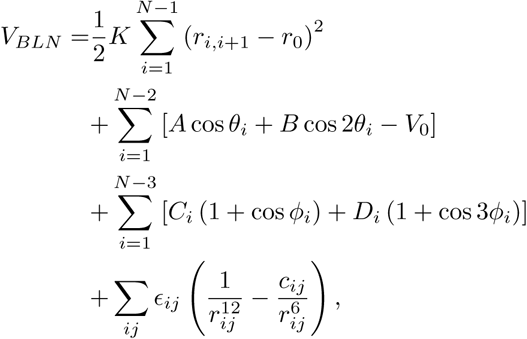

where *r*_*ij*_ is the distance between residues *i* and *j, θ*_*i*_ is the angle between three contiguous residues, and *φ* the dihedral angle. The sequence-independent parameters for the model are *K* = 50, *r*_0_ = 1, *A* = 5.118, *B* = 5.308, *V*_0_ = − 5.295, while the sequence-dependent: *C*_*i*_ = 0 and *D*_*i*_ = 0.2 if two or more residues are neutral, or *C*_*i*_ = *D*_*i*_ = 1.2 otherwise; while the Lennard-Jones: *c*_*ij*_ = 0, *E*_*ij*_ = 4 if either *i* or *j* are neutral, *c*_*ij*_ = 1, and *E*_*ij*_ = 4 if *i* and *j* are hydrophobic, and *c*_*ij*_ = − 1, and *E* = 8*/*3 otherwise, all adimensional units.

Physical units are estimated in the following way. Length units are recovered assuming that the distance between *α* carbons in a protein is 0.38 nm. The energy units are defined assuming the energy of a hydrogen bond *ϵ* ≈ 1.7*kT*, which defines force units as 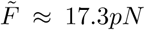. Mass units assume the average mass of an aminoacid *m*_*a*_ ≈ 3 *×* 10^−22^kg. Thus, time units can be defined, as: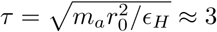 ps.

We carry out all simulations using a self-written code, integrating the overdamped Langevin equations using a stochastic second order Runge-Kutta algorithm^55^. The integration step is Δ*t* = 0.005*τ*.

